# Gibberellin-deactivating GA2OX enzymes act as a hub for auxin-gibberellin crosstalk in *Arabidopsis thaliana* root growth regulation

**DOI:** 10.1101/2025.02.03.636207

**Authors:** Monika Kubalová, Jayne Griffiths, Karel Müller, Alexander M. Jones, Matyáš Fendrych

## Abstract

Plant bodies are built from immobile cells, making the regulation of cell expansion essential for growth, development, and adaptation. In roots, cell elongation executes the movement of the root tips through soil. This process is tightly controlled by numerous signaling pathways. Among these, gibberellin and auxin signaling stand out for their contrasting effects on root growth, interacting through complex crosstalk at multiple regulatory levels. Here we reveal the molecular basis of the auxin-gibberellin crosstalk in the model plant *Arabidopsis thaliana*. We show that auxin signaling pathway steers the expression of *GIBBERELLIN 2-OXIDASES (GA2OX)*, key gibberellin-deactivating enzymes in the root elongation zone. GA2OX are negative regulators of root cell elongation; *GA2OX8* overexpression decreases gibberellin levels and inhibits root cell elongation, in contrast, the *ga2ox heptuple* mutant roots show elevated gibberellin levels in the elongation zone and grow longer roots. Shoot derived auxin can regulate *GA2OX8* expression in roots, linking systemic auxin signaling to local gibberellin modulation. In addition, *GA2OX8* is active in vascular tissues and the stem cell niche, tissues with high auxin levels. Loss of *GA2OX* genes results in altered stem cell niche, including increased quiescent center size and expanded root cell layers, highlighting the role of these enzymes in maintaining tissue organization. Together, our findings identify GA2OX6 and GA2OX8 enzymes as key mediators of auxin-gibberellin crosstalk, providing insights into their roles in root elongation, vascular development, and stem cell niche maintenance. These results expand our understanding of how auxin integrates with gibberellin signaling to coordinate root development and growth dynamics.

## Introduction

Plants show a remarkable ability to modulate growth in response to developmental and environmental cues. The activity of root meristem and elongation zone is orchestrated by a complex network of phytohormonal signalling pathways to steer the navigation of the root through the soil, and to ensure coordinated growth of aerial organs with roots. The phytohormones auxin and gibberellin (GA) play pivotal and sometimes opposing roles in the control of cellular elongation in roots.

The nuclear auxin pathway (NAP) is characterized by the auxin receptor TRANSPORT INHIBITOR RESPONSE1 (TIR1)/AUXIN-SIGNALLING F-BOX (AFB), which, in the presence of auxin, associates with coreceptors AUXIN/INDOLE ACETIC ACID (Aux/IAA) proteins, leading to their degradation and the release of AUXIN RESPONSE FACTOR (ARF) that modulate the transcription of auxin-regulated gene^1^. As far back as 1939, it was discovered that disturbing NAP inhibits root elongation^2^. Surprisingly, recent study shows that inhibiting NAP initially boosts root cell elongation, and only after several hours, it leads to meristem exhaustion, resulting in well-known inhibition of root growth^3^. This highlights the time dependency of auxin responses in the root. Additionally, a non-transcriptional auxin signalling pathway functions in *Arabidopsis thaliana* roots and appears to interact with the transcriptional pathway^3–8^.

GA are master regulators of cellular elongation^9^. GA bind their receptor GIBBERELLIN INSENSITIVE DWARF (GIDs) and DELLA transcriptional repressors (including RGA and GAI), leading to the degradation of DELLA and subsequent activation of GA-regulated gene transcription^10–13^. The *Arabidopsis thaliana* root tip exhibits a remarkable gradient of bioactive GA concentration along the primary root axis, rising with distance from the meristem and peaking in the elongation zone^3,14,15^, which is consistent with GA being positive drivers of cellular elongation. Additionally, GA regulate the cell cycle^16^ and control axial root development^17,18^. The spatial and temporal distribution of GA is determined by a combination of biosynthetic, catabolic, and transport processes^19^. To produce bioactive GA, most common precursors GA12 or GA53 undergo a series of reactions mediated by the group of biosynthetic enzymes – GIBBERELLIN OXIDASES - GA20OX and GA3OX^20^. Inactivation of bioactive GA is catalysed by GA2OX enzymes^21^. The GA2OX enzyme family in *Arabidopsis thaliana* consists of nine members: *GA2OX1, GA2OX2, GA2OX3, GA2OX4, GA2OX6, GA2OX7, GA2OX8, GA2OX9* and *GA2OX10*^21^. Overexpression of *GA2OX8* and *GA2OX7*^22,23^ or *GA2OX9* and *GA2OX10*^24^ genes leads to reduced levels of bioactive GA and results in dwarfism, delayed flowering, and changed leaf morphology. On the other hand, higher order loss of function *ga2ox* mutants show elevated levels of active GA compared to the wild type. Consequently, they display phenotypes indicative of GA excess^23–25^. Although the importance of GA2OX in aboveground tissue development is clear, their role in root development is still not understood.

Auxin and GA signalling pathways are interconnected and interact with each other, either positively or negatively during various developmental responses^26^. This has been studied at different levels, including their mutual effect on signalling, deactivation^27–29^, transport^28,30,31^, and biosynthesis^27,32–34^.

In Arabidopsis root, GA modulate auxin transport to enhance root responsiveness to auxin^28^ or to regulate vascular development^35^. Additionally, shoot-derived auxin modulates root growth by regulating the effect of GA response^29^. Initial inhibition of NAP increases sensitivity of root to GA^3^. Although GA and auxin clearly affect each other during root development, how auxin and GA actions overlap during root cell elongation is not completely understood.

Here, to understand the nature and physiological effects of auxin-GA crosstalk during root cell elongation, we focus on two auxin-regulated GA-deactivating enzymes GA2OX6 and GA2OX8. We show that NAP components IAA17/AXR3 and ARF5/MP regulate the expression of *GA2OX6* and *GA2OX8* in the root elongation zone. Consequently, GA2OX6 and GA2OX8 decrease the level of GA in the elongation zone, contributing to the regulation of root cell elongation. Our findings reveal the genetic and molecular mechanisms by which the phytohormone auxin regulates the levels of bioactive GA in the root elongation zone of *Arabidopsis thaliana*.

## Results

### Auxin decreases GA response and GA level in Arabidopsis root tissue

We have shown previously that by a genetic manipulation of the nuclear auxin pathway (NAP) we can steer the elongation rate of Arabidopsis roots; the inducible overexpression of a dominant inhibitor (dominant *IAA17/AXR3 -* pG1090::XVE>>*AXR3-1*) or activator (dominant *ARF5/MP –* pG1090::XVE>>*ΔMP*) of auxin-induced transcription leads to root growth promotion and growth inhibition, respectively^3^. We harnessed this system to obtain a time-resolved transcriptomic profiles of the roots with altered cellular elongation rates (Fig.1A).

**Figure 1.**
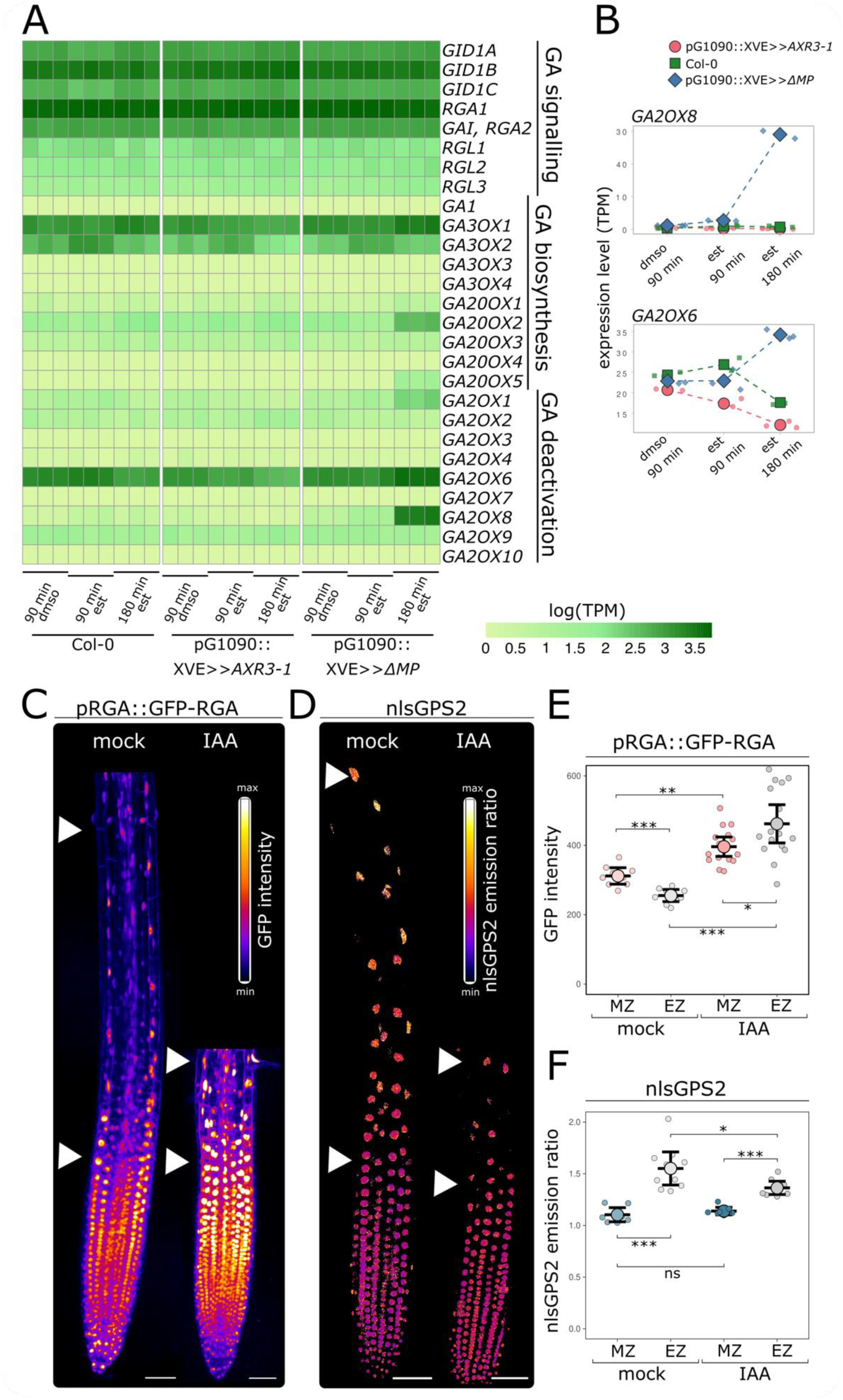
IAA treatment results in a decrease in both the GA response and GA levels in Arabidopsis root tissues. A. Heat map of GA related genes generated from RNAseq data reflecting Log-transformed TPM (transcripts per million) values in Col-0, pG1090::XVE>>*AXR3-1*and pG1090::XVE>>*ΔMP* roots after indicated treatments. B. Relative expression level of *GA2OX6* and *GA2OX8* represented as TPM in pG1090::XVE>>*AXR3-1*, pG1090::XVE>>*ΔMP* and Col-0 root cells after 90min of dmso or 90 or 180min of estradiol treatment. C. Root tip of pRGA::GFP-RGA treated with 50nM IAA or mock for 5h. White arrows indicate the beginning and the end of the elongation zone. Scale bar = 50μM. D. Root tip of nlsGPS2 treated with 50nM IAA or mock for 5h. White arrows indicate the beginning and the end of the elongation zone. Scale bar = 50μM. E. Quantification of GFP-RGA represented as 488 signal intensity of nuclei in meristematic zone (MZ) and elongation zone (EZ) of mock or IAA treated pRGA::GFP-RGA roots. F. Quantification of nlsGPS2 emission ratio represented as FRET/458 ratio of nuclei in meristematic zone (MZ) and elongation zone (EZ) of mock or IAA treated nlsGPS2 roots. The asterisks indicate statistically significant differences (*P < 0.05, **P < 0.01, ***P < 0.001). Error bars in boxplots are CI.

Among the differentially expressed genes, we noticed a conspicuous correlation between NAP activity and GA-deactivating genes *GA2OX1, GA2OX6* and *GA2OX8*. *GA2OX6* and *GA2OX8* showed the strongest auxin signalling-dependent induction among the *GA2OX* family genes. Activation of auxin signalling by the *ΔMP* strongly upregulated both genes while inhibition of the pathway by *AXR3-1* induction led to their downregulation. *GA2OX6* is the strongly expressed in the root and *GA2OX8* shows the strongest upregulation upon activation of NAP (Fig.1A,B). Additionally, the rapid response of *GA2OX6* and *GA2OX8* to auxin treatment in root tip^3^ primed them for selection as candidate genes. Two of the GA biosynthetic genes showed a mild upregulation upon activation of auxin signalling but were unchanged in the induced *AXR3-1* roots. The GA signalling genes showed no significant changes (Fig.1A, Dataset1). This indicates that auxin signalling could lower the levels of active GA in Arabidopsis root tips via activation of GA2OX enzymes that catabolise bioactive GA and its precursors.

Therefore, we analyzed the spatial and temporal dynamics of GA signalling (Fig.1C) and levels of bioactive GA in roots (Fig.1D) upon auxin treatment. We utilized the GA signalling marker line pRGA::GFP-RGA^12^ in which the GFP signal intensity decreases with activation of GA signalling. To visualize the levels of GA in roots, we used the direct FRET-based nuclear localised nlsGPS2 (GIBBERELLIN PERCEPTION SENSOR 2) biosensor that was recently engineered to be more orthogonal and reversible^36^. Both reporter and biosensor lines showed increased GA response (Fig.1C,E) or GA level (Fig.1D,F) in root elongation zone compared to the meristematic zone, corresponding to the published data^3,14,15^. A 5h treatment of roots with naturally occurring auxin - Indole-3-acetic acid (IAA) led to a significant increase of pRGA::GFP-RGA signal intensity in both meristematic and elongation zones (Fig.1C,E), indicating decreased GA signalling after IAA treatment. The nlsGPS2 biosensor emission ratio decreased after IAA treatment (Fig.1D,F), indicating a decrease in cellular GA level. Both genetic markers thus show that auxin treatment causes a decrease in GA levels and GA signalling response in the elongation zone of Arabidopsis roots. The plausible explanation is that this effect is mediated by the activation of *GA2OX6* and *GA2OX8* genes.

### Auxin signalling steers the expression of GA deactivating genes *GA2OX6* and *GA2OX8* in the root elongation zone

The analysis of *GA2OX6* and *GA2OX8* promoters revealed the presence of AuxRE ARF binding sites (Fig.S1A). To test whether auxin signalling regulates the activity of the *GA2OX* genes in planta, we first used the transient *Nicotiana benthamiana* expression system to score the effect of *AXR3-1* on the activity of the *GA2OX8* promoter. *Luciferase* gene (*LUC*) driven by *GA2OX8* promoter was co-expressed with the dominant inhibitor *AXR3-1* tagged with mVenus and/or with wild type form of *ARF5/MP* tagged with mScarlet (Fig.2A). The expression of the constructs was verified using confocal microscopy (Fig.S1B). The co-expression with *AXR3-1* inhibited the activity of *GA2OX8* promoter (Fig.2 A,B). *AXR3-1* action was not dependent on the expression of *ARF5/MP* (Fig.2 A,B), which can be explained by binding of the tobacco ARFs to the AuxRE in the *GA2OX8* promoter.

**Figure 2:**
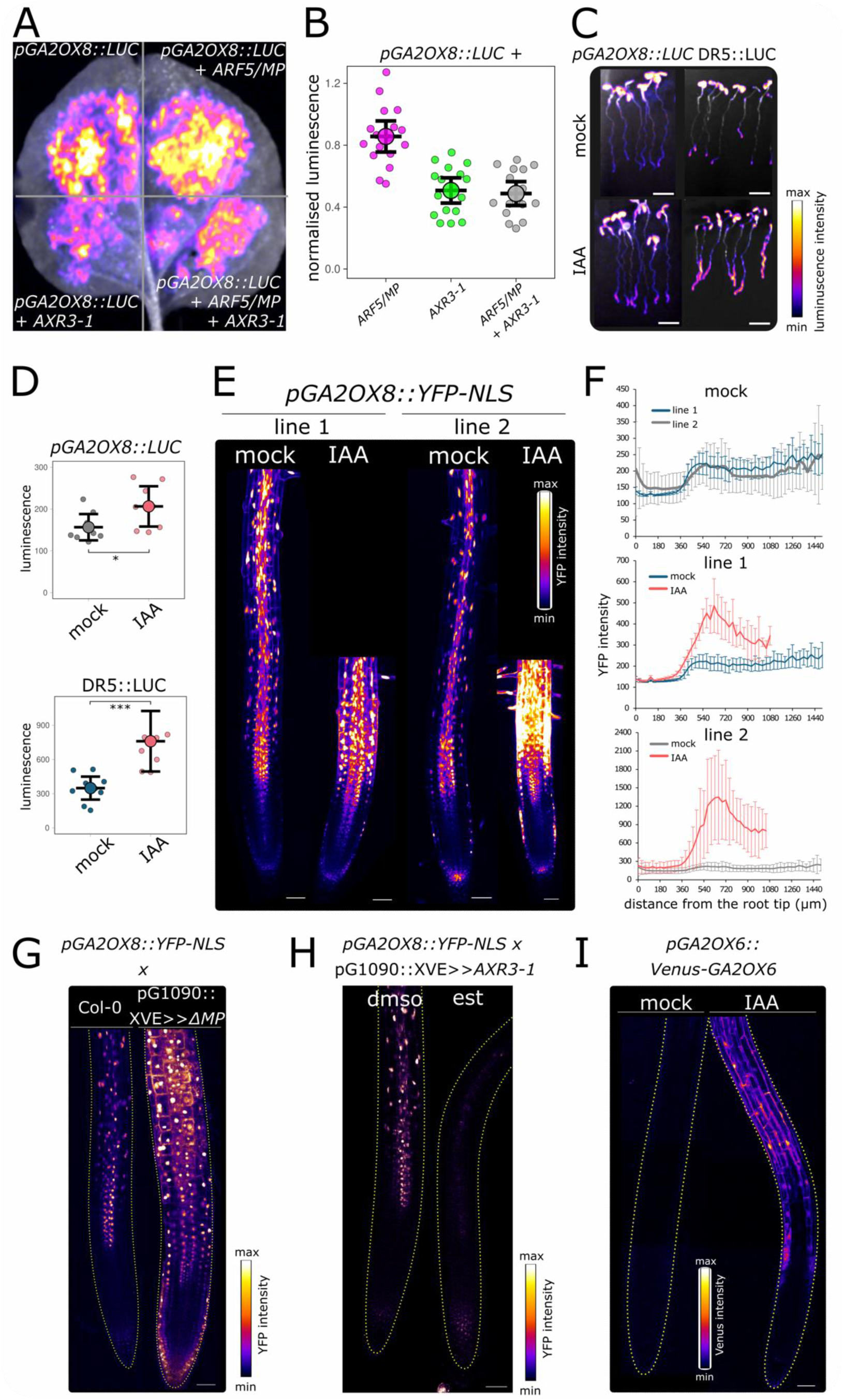
GA-deactivating enzymes GA2OX6/8 are regulated by auxin-IAA17/AXR3-ARF5/MP pathway. A. Activity *pGA2OX8::LUC* in tabacco leaf co-infiltrated with/without p35S::*ARF5-mScarlet* and p35S::*AXR3-1-mVenus*. B. Quantification of *pGA2OX8::LUC* activity co-infiltrated with/without p35S::*ARF5-mScarlet* and p35S::*AXR3-1-mVenus*. Normalised to luminescence of *pGA2OX8::LUC* without co-infiltration. C. Activity of luciferase in DR5::LUC and *pGA2OX8::LUC* plants treated with mock or 50nM IAA for 5h. Scale bar = 5mm. D. Quantification of luciferase activity in root tips of DR5::LUC and *pGA2OX8::LUC* plants treated with mock or 50nM IAA for 5h. E. Expression of *pGA2OX8::YFP-NLS* of 2 independent homozygous lines in root tip treated with mock or 50nM IAA for 5h, represented as 515 signal intensity. Scale bar = 50μM. F. Quantification of YFP-NLS represented as 515 signal intensity along the longitudinal axis of the root in mock or IAA treated *pGA2OX8::YFP-NLS* lines. G. Expression of YFP-NLS in root tips of *pGA2OX8::YFP-NLS* crossed with pG1090::XVE>>*ΔMP* or Col-0, treated for 24h with estradiol. The outlines of the root are highlighted with a yellow dashed line. Scale bar = 50μM. H. Expression of YFP-NLS in root tips of *pGA2OX8::YFP-NLS* crossed with pG1090::XVE>>*AXR3-1* treated for 24h with dmso or estradiol. The outlines of the root are highlighted with a yellow dashed line. Scale bar = 50μM. I. Expression of *pGA2OX6::Venus-GA2OX6* in root tip treated with mock or 10nM IAA for 90min, represented as 515 signal intensity. The outlines of the root are highlighted with a yellow dashed line. Scale bar = 50μM. The asterisks indicate statistically significant differences (*P < 0.05, **P < 0.01, ***P < 0.001). Error bars in boxplots are CI and in line graph are SD.

Next, to monitor the activity of the *GA2OX8* promoter on the organ level, we introduced *pGA2OX8::LUC* to Arabidopsis plants, and we observed a strong expression in roots and cotyledons of seedlings. The treatment with IAA led to an increase in LUC luminescence, which was particularly apparent in the younger parts of roots (Fig.2C,D). Similarly, the positive control – *LUC* gene driven by the synthetic *DR5* promoter^37^ showed luminescence increase in response to the IAA treatment. Further, we prepared a transcriptional *GA2OX8* promoter reporter line driving the expression of nuclear-localized YFP (*pGA2OX8::YFP-NLS*, Fig.2E), which allows for the analysis of promoter activity on the cellular level. In the root tip, we could detect weak *pGA2OX8::YFP-NLS* signal in the lateral root cap cells and the stele tissues of the meristematic zone. A strong expression started in the elongation zone of the root where the signal was visible both in the stele tissues and the outer tissues epidermis and cortex, and the expression remained high in the root hair zone (Fig.2E,F). The treatment with IAA dramatically increased the *GA2OX8* promoter activity (Fig.2E,F). Notably, this increase of *GA2OX8* promoter activity was visible specifically in the elongation zone of the root, implying its involvement in the regulation of root cell elongation.

To confirm direct regulation of *GA2OX8* promoter by NAP components, we crossed *pGA2OX8::YFP-NLS* line with the inducible dominant pG1090::XVE>>*AXR3-1* and pG1090::XVE>>*ΔMP* lines that inhibit and stimulate auxin signalling, respectively. Induction of irrepressible *ΔMP* led to a significant increase of *GA2OX8* promoter activity in most root tip tissues (Fig.2G). In contrast, the induction of *AXR3-1* almost completely abolished activity of *GA2OX8* promoter in all root tip tissues (Fig.2H).

Finally, we took the advantage of the published *GA2OX6* translational reporter line (*pGA2OX6::Venus-GA2OX6*^38^) to analyze the localization of this gene expression and monitor the fusion protein levels in the roots. The Venus-GA2OX6 protein fluorescence was not detectable in the root tips under normal growth conditions. However, already after 90min of IAA treatment, the Venus-GA2OX6 signal was clearly detectable in the epidermis of the elongation and maturation zone of the root (Fig.2I).

Altogether, these results clearly show that auxin, through NAP’s components IAA17/AXR3 and ARF5/MP, steers the expression levels of *GA2OX8/6* genes in roots and the elongation zone in particular.

### *GA2OX* genes steer root cell elongation

To study the role of GA2OX enzymes in root growth regulation, we analysed root phenotypes of plants with altered expression of *GA2OXs* genes. First, we prepared plants expressing *GA2OX8* from a strong estradiol-inducible system - pG1090::XVE>>*mCherry-GA2OX8* (Fig.3A). The induction of *GA2OX8* led to decrease of GA level (Fig.3B,C) and GA response (Fig.3D,E) in both the meristem and the elongation zone. This indicates that the *mCherry-GA2OX8* fusion protein is functional and decreases the level of bioactive GA.

**Figure 3:**
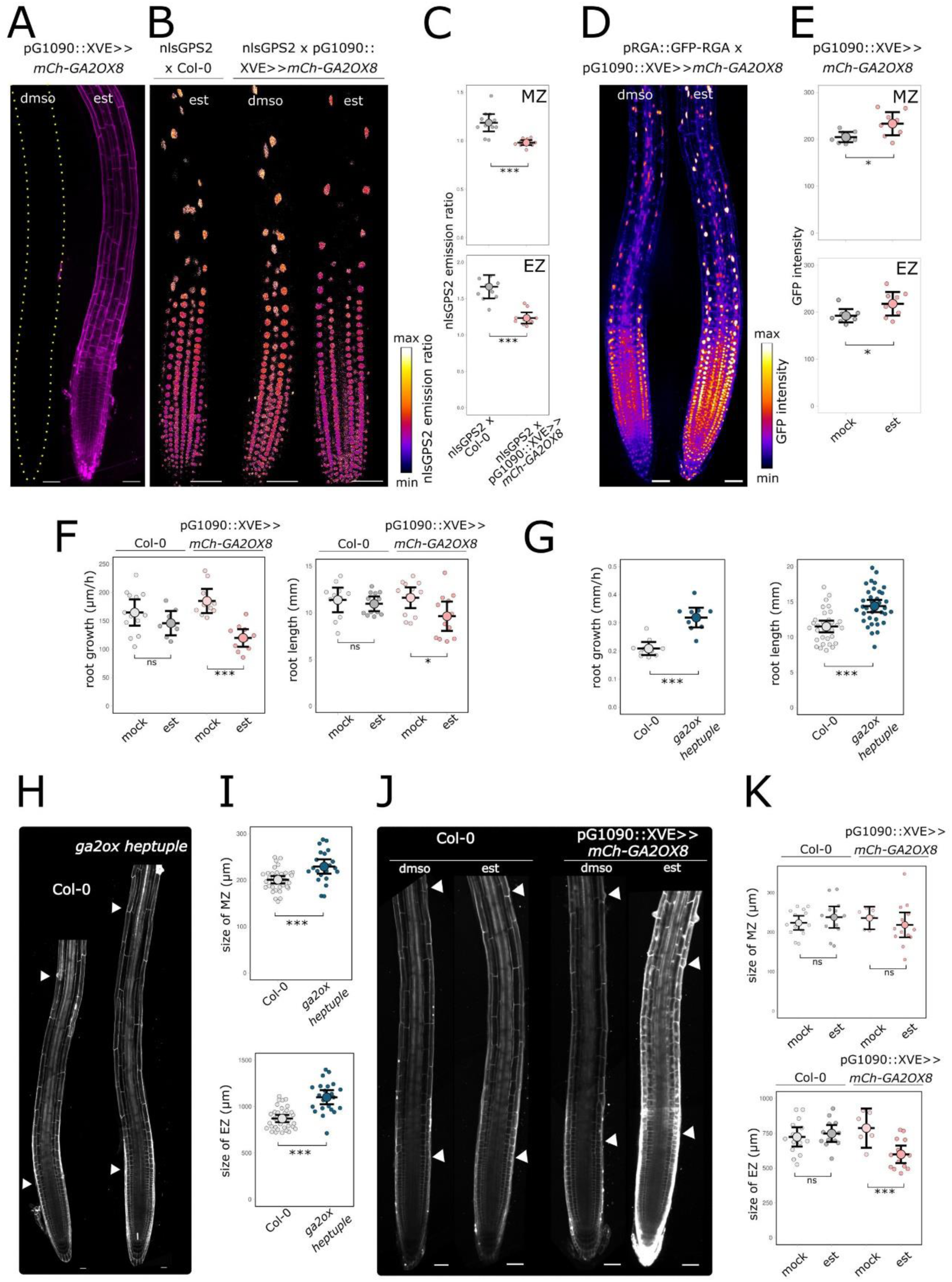
GA2OXs enzymes regulate root cell elongation. A. Root tip of pG1090::XVE>>*mCherry-GA2OX8* treated 24h with estradiol or dmso. The outlines of the root are highlighted with a yellow dashed line. Scale bar = 50μM. B. Root tips of Col-0 x nlsGPS2 and pG1090::XVE>>*mCherry-GA2OX8* x nlsGPS2 treated for 24h with estradiol or dmso. Scale bar = 50μM. C. Quantification of nlsGPS2 emission ratio (FRET/458 ratio) corresponding to GA level of nuclei in meristematic (MZ) and elongation zone (EZ) of pG1090::XVE>>*mCherry-GA2OX8* x nlsGPS2 treated for 24h with estradiol or dmso. D. Root tips of pG1090::XVE>>*mCherry-GA2OX8* x pRGA::GFP-RGA treated for 24h with estradiol or dmso. Scale bar = 50μM. E. Quantification of the GFP-RGA intensity represented as 488 signal intensity of nuclei in meristematic (MZ) and elongation zone (EZ) of pG1090::XVE>>*mCherry-GA2OX8* x pRGA::GFP-RGA treated for 24h with estradiol or dmso. F. Quantification of root length of pG1090::XVE>>*mCherry-GA2OX8* and Col-0 grown on estradiol and growth rate of pG1090::XVE>>*mCherry-GA2OX8* and Col-0 treated 24h with estradiol. G. Quantification of root length and root growth rate of *ga2ox heptuple* mutant and Col-0. H. Root tips of *ga2ox heptuple* mutant and Col-0 stained with propidium iodide (PI). White arrows indicate the beginning and the end of the elongation zone. Scale bar = 50μM. I. Quantification of the size of meristematic zone (MZ) and elongation zone (EZ) of *ga2ox heptuple* mutant and Col-0. J. Root tips of pG1090::XVE>>*mCherry-GA2OX8* and Col-0 treated for 24h with estradiol or mock stained with PI. White arrows indicate the beginning and the end of the elongation zone. Signal in estradiol treated pG1090::XVE>>*mCherry-GA2OX8* is the combination of PI and mCherry intensity. Scale bar = 50μM. K. Quantification of the size of meristematic zone (MZ) and elongation zone (EZ) of pG1090::XVE>>*mCherry-GA2OX8* and Col-0 treated for 24h with estradiol or mock. The asterisks indicate statistically significant differences (*P < 0.05, **P < 0.01, ***P < 0.001). Error bars in boxplots are CI.

To see the direct effect of *GA2OX8* expression on root growth, we measured root growth rate 24h after *GA2OX8* induction. Overexpression and accumulation of GA2OX8 led to shorter, slower growing roots (Fig.3F).

Second, to assess physiological importance of the root-expressed *GA2OX* genes, we analyzed a series of loss-of-function mutants. Firstly, we tested 2 candidates: *GA2OX6* which is strongly expressed in the root and *GA2OX8* which shows the strongest upregulation upon activation of NAP. Root length of single *ga2ox6* or *ga2ox8* T-DNA insertion mutant did not show any significant differences compared to control, the same was true for the *ga2ox6xga2ox8 double* mutant (Fig.S1C). As there are 9 characterised *GA2OX* genes in Arabidopsis^21^, this can be the result of gene redundancy. Therefore, we prepared a mutant in seven *GA2OX1,2,3,4,6,7,8* genes (*ga2ox heptuple* mutant) and analysed its root phenotype. Mutation in seven *GA2OX* genes led to longer and faster growing roots (Fig.3G). To understand the physiological changes caused by altered level of *GA2OX* genes, we analysed the longitudinal zonation of pG1090::XVE>>*mCherry-GA2OX8* and *ga2ox heptuple* mutant. While mutation of *GA2OX* genes resulted in longer elongation zone (Fig.3H,I), overexpression of *GA2OX8* gene resulted in shorter elongation zone (Fig.3J,K). Interestingly, *ga2ox heptuple* mutant has a slightly longer meristematic zone (Fig.3I). However, overexpression of *GA2OX8* did not induce the opposite alteration (Fig.3K).

To sum up, these results show that GA2OX8 protein decreases the level of bioactive GA in the roots and that this way it regulates root growth predominantly by decreasing the root elongation zone size.

### Auxin regulates the level of GA in root elongation zone through *GA2OX*s

The regulation of GA deactivating genes *GA2OX6* and *GA2OX8* by auxin represents a crosstalk between these two hormonal pathways. We wanted to address the physiological significance of this crosstalk in Arabidopsis roots. First, to test whether auxin directly regulates GA level through *GA2OX* genes, we analysed GA level in *ga2ox heptuple* mutant after IAA treatment. For this, we transformed *ga2ox heptuple* mutant with nlsGPS2 sensor (Fig.4A). As expected, the *ga2ox heptuple* showed significantly increased level of GA in the root elongation zone, while GA level in the meristematic zone was comparable to control (Fig.4 A,B,C). IAA treatment did not cause any changes of GA level in meristematic zone, regardless of genotype (Fig.4 A,B,C). In contrast, in response to IAA, GA level decreased in the elongation zone of the control plants, while the *ga2ox heptuple* roots showed a partial resistance to IAA; in the mutant, the nlsGPS2 emission ratio decrease was smaller than in the control (Fig.4 A,C). This suggests that auxin signalling regulates GA level in the root elongation zone through *GA2OX* genes.

**Figure 4:**
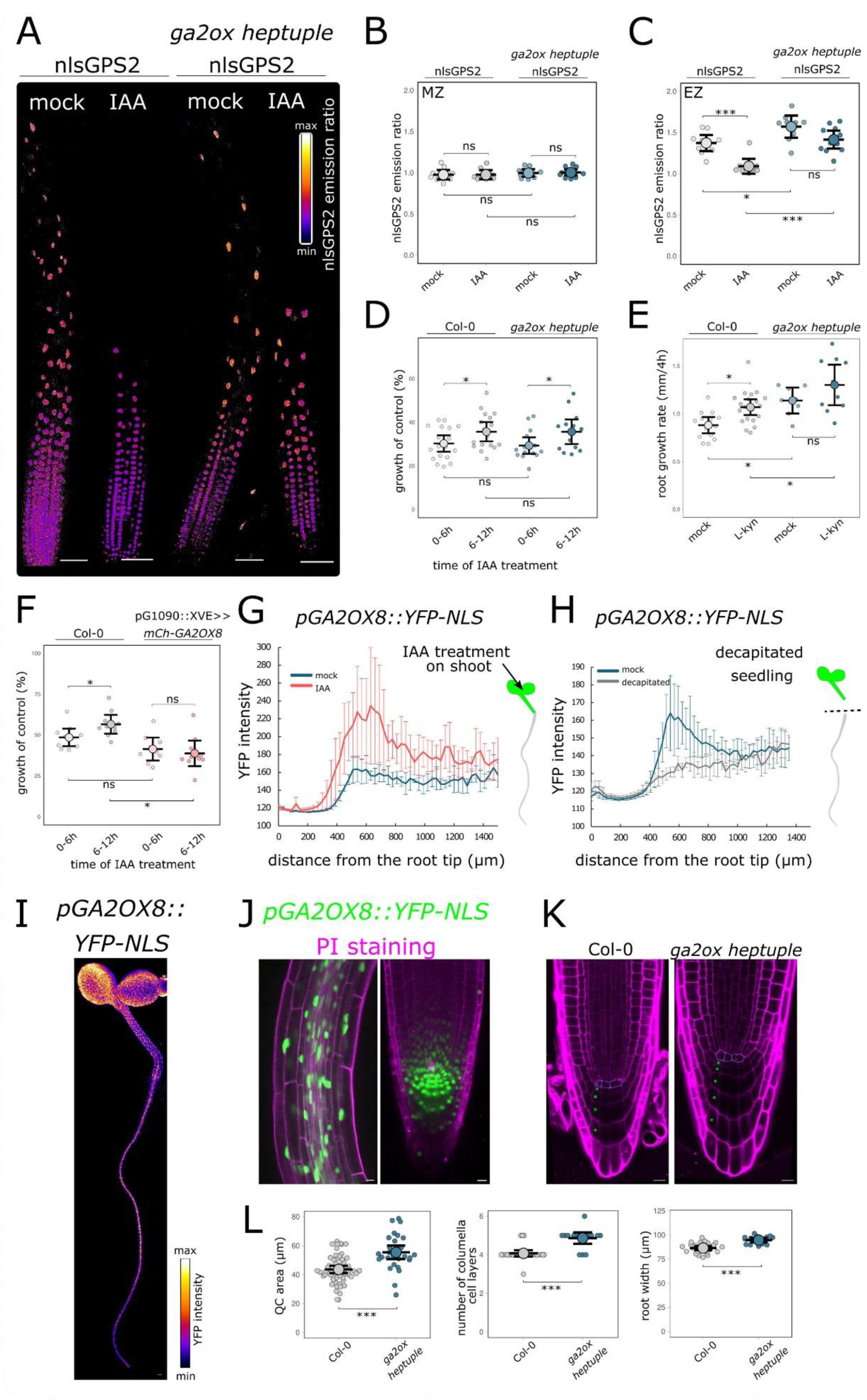
Auxin signaling directly regulates GA2OX genes. A. Root tips of nlsGPS2 and *ga2ox heptuple* nlsGPS2 treated for 5h with 50nM IAA or mock. Scale bar = 50μM. B. Quantification of nlsGPS2 emission ratio (FRET/458 ratio) corresponding to GA level of nuclei in the meristematic zone (MZ) of nlsGPS2 and *ga2ox heptuple* nlsGPS2 treated for 5h with 50nM IAA or mock. C. Quantification of nlsGPS2 emission ratio (FRET/458 ratio) corresponding to GA level of nuclei in the elongation zone (EZ) of nlsGPS2 and *ga2ox heptuple* nlsGPS2 treated for 5h with 50nM IAA or mock. D. Growth rate of *ga2ox heptuple* and Col-0 treated with 5nM IAA in 0-6h (initial phase) and 6-12h (recovery phase) after treatment. Normalized to the mock-treated growth rate. E. Growth rate of *ga2ox heptuple* and Col-0 treated with 1.5μM L-Kyn (inhibitor of auxin biosynthesis) for 4h. F. Growth rate of pG1090::XVE>>*mCherry-GA2OX8* and Col-0 treated with 5nM IAA and estradiol in 0-6h (initial phase) and 6-12h (recovery phase) after treatment. Normalized to the mock-treated growth rate. pG1090::XVE>>*mCherry-GA2OX8* and Col-0 grown on estradiol. G. Quantification of YFP-NLS represented as 515 signal intensity along the longitudinal axis of *pGA2OX8::YFP-NLS* roots in plants treated for 5h with 100μM IAA/mock on shoot. H. Quantification of YFP-NLS represented as 515 signal intensity along the longitudinal axis of *pGA2OX8::YFP-NLS* roots 16h after decapitation. I. Expression of *pGA2OX8::YFP-NLS* in Arabidopsis plant. Scale bar = 100μM. J. Expression of *pGA2OX8::YFP-NLS* (green) in root vascular tissue (left) and root tip (right) stained with PI (magenta). Scale bar = 50μM. K. Root tips of *ga2ox heptuple* and Col-0 stained with PI (magenta). The outlines of the QC cells are highlighted with a blue dashed line, green asterisks indicate columella cell layers. Scale bar = 10μM. L. Quantification of QC area, number of columella cell layers and root width of *ga2ox heptuple* and Col-0. The asterisks indicate statistically significant differences (*P < 0.05, **P < 0.01, ***P < 0.001). Error bars in boxplots are CI and in line graph are SD.

Further, we tested whether this crosstalk can affect the root growth response to auxin. To this end, we treated *ga2ox heptuple* and pG1090::XVE>>*mCherry-GA2OX8* line with low concentration of IAA and measured the root growth inhibition rate after 0-6 and 6-12 hours. After initial growth inhibition, control and *ga2ox heptuple* roots partially recovered from growth inhibition (Fig.4D). Interestingly, *ga2ox heptuple* mutant was less sensitive to increased root cell elongation caused by inhibition of auxin biosynthesis (Fig.4E). Additionally, root growth of plants overexpressing *GA2OX8* is significantly more inhibited after longer IAA treatment (6-12h), indicating that they are hypersensitive to auxin (Fig.4F).

It was shown that shoot-derived auxin is essential for GA signalling during root growth^29^. Is *GA2OX8* a molecular link between shoot-derived auxin and regulation of GA signalling in root? To test whether auxin arriving from the shoot drives the *GA2OX8* expression in root, we applied exogenous IAA to cotyledons. On the other hand, to decrease the flux of auxin from shoot to root, we decapitated the seedlings. IAA application to cotyledons led to increase of *GA2OX8* expression in root tissue (Fig.4G) and removing shoot as a source of auxin resulted in decreased *GA2OX8* expression predominantly in root elongation zone (Fig.4H), showing that shoot-derived auxin regulates *GA2OX8* gene expression in root elongation zone.

These results suggest that GA2OX-modulated GA level contributes to regulation of root cell elongation in response to auxin, including shoot derived auxin. Although *GA2OX* genes are included in auxin-mediated regulation of root cell elongation, they do not seem to be essential.

### *GA2OX8* is expressed in vascular tissue and stem cell niche

NAP positively regulates the expression of *GA2OX6* and *GA2OX8.* We aimed to investigate whether this mechanism plays a role in other biological processes beyond root cell elongation. To answer this question, we analysed activity of *GA2OX8* promoter within the plant.

*GA2OX8* promoter is active in all plant tissues (Fig.4I), we focus on its expression in auxin-rich tissues. Particularly interesting is the strong expression of *GA2OX8* in the root vascular system, both in xylem and phloem cells (Fig.4J). GA2OX8 may be involved in regulation of appropriate GA level in stele necessary for vascular development^35^.

Additionally, *GA2OX8* gene is expressed in stem cell niche (SCN), including quiescent centre (QC) (Fig.4J), corresponding to auxin maxima in these cells^39^. QC is a signalling hub that maintains the surrounding stem cells in an undifferentiated state regulating the balance between cell division and differentiation in the root tip. To test whether *GA2OXs* are involved in this process, we focused on the architecture of root tip in *ga2ox heptuple* mutant (Fig.4K). Plants lacking *GA2OX* genes showed bigger QC, more columella cell layers and wider root cell layers (Fig.4K,L).

These results suggest that GA2OX8 may be involved in elimination of excessive GA in SCN to influence cell division rates, cell expansion or cell fate.

Altogether, we show that GA2OX enzymes are negative regulators of root cell elongation and represent a hub for the crosstalk between auxin and GA signalling during root cell elongation, vascular development and regulation of stem cell niche.

## Discussion

Highly coordinated hormonal regulation is fundamental for root cell elongation^40^. This study shows the role of auxin-modulated GA deactivating enzymes GA2OX in root growth and development. An increase in GA level positively correlates with root cell elongation rate^3,14,15^. In this work we show that auxin treatment decreases GA level and alters the shape of GA concentration gradient along the longitudinal axis of the root. Auxin-treatment thus results in inhibited root cell elongation and a perturbation of GA gradient in root.

We showed that GA deactivating enzymes GA2OX6 and GA2OX8 participate in the auxin-induced GA decrease in elongation zone of Arabidopsis root. Previously, it has been shown that these genes are specifically regulated by auxin at the whole plant level^27^. In roots, these genes are upregulated already 20min after IAA treatment^3^. GA2OX6 and GA2OX8 reduced growth of aboveground parts^22,23,41^, however their role in roots has not been characterized. Here, we demonstrate that GA2OX6 and GA2OX8 enzymes deactivate GA and are upregulated by auxin treatment in the elongation zone of the Arabidopsis root. Considering that there is no characterized GA exporter, GA2OX activity can act as main negative regulators of GA levels in the root elongation zone. Remarkably, the *ga2ox septuple* mutant showed an increase in GA levels specifically in the root elongation zone, while GA levels in the meristem remained unchanged. The exact same conclusion was reached by Kiradjiev (unpublished), confirming that GA2OX enzymes are crucial determinants of the GA gradient in the root.

What is the biological role of GA2OXenzymes in root? Both the mutant and the *GA2OX* overexpression data demonstrate that GA2OXs are negative regulators of root cell elongation. The absence of a phenotype in single or double *ga2ox* mutants points to gene redundancy, which is remarkable for GA metabolic genes, as overlapping functions have been demonstrated even among genes that show non-overlapping expression pattern under normal conditions^42^. Even the mutation of 5 *GA2OX* genes did not affect GA levels, which corresponds to the unchanged root length and elongation growth of *ga2ox quintuple* mutant^43^. Using the inducible system, we were able to study the direct effect of *GA2OX8* gene induction on root growth while excluding the possible influence of GA level changes on other processes, such as seed germination rate. Additionally, germination is likely regulated by de novo GA biosynthesis, as indicated by the unchanged GA level in *ga2ox quintuple* mutant seeds^25^.

The primary driver of GA levels in root elongation zone is the activity of biosynthetic enzymes and the delivery of GA precursors via phloem (Kiradjiev et al., unpublished). In addition to differential GA biosynthesis and cellular permeability to GA^43^, the deactivation by GA2OX genes can jointly regulate GA levels in the elongation zone. Interestingly, ectopic overexpression of biosynthetic genes and the subsequent increase in GA levels did not lead to enhanced elongation in the early elongation zone but rather in later tissues^43^. This suggests that spatially specific effects of mutating *GA2OX* genes contribute to the regulation of elongation growth.

GA2OX6 and GA2OX8 enzymes were shown to degrade the biologically active GA4 and the GA4 precursor, respectively^23,65^. Based on a multiscale model of GA gradient formation in roots, Kiradjiev et al (unpublished) propose that degradation of bioactive GA will have a stronger impact on GA levels than degradation of GA precursors. Our data show that overexpression of GA2OX8 significantly affects GA levels and growth of the roots. Similarly, the levels of bioactive GA4 were decreased in the GA2OX8 overexpressing lines. Additionally, Kiradjiev et al (unpublished) postulate that phloem unloading of the shoot-derived GA precursors is crucial for the level of bioactive GA in the elongation zone. It is possible that, apart from the local degradation of GA precursors in the root, the delivery of GA precursors is decreased in our GA2OX8 overexpressor line.

*GA2OX6* and *GA2OX8* are downstream targets of auxin signalling, representing a crosstalk between these phytohormonal pathways. What is the biological significance of this crosstalk for root growth regulation? In the *ga2ox heptuple* mutant, the IAA-triggered decrease of GA level was less significant than in the control. This “molecular phenotype” demonstrates that GA2OX6 and GA2OX8 decrease GA level in the root elongation zone in response to increased auxin input. Accordingly, at lowered auxin level, *ga2ox heptuple* mutant did not significantly increase elongation rate. This suggests that this crosstalk might reinforce the negative auxin effect on cell elongation. In agreement with this hypothesis, the *GA2OX8* overexpressing plants were mildly hypersensitive to external auxin application. However, the control-like sensitivity of *ga2ox heptuple* root to exogenous auxin treatment contradicts this hypothesis. A possible explanation could be the *GA2OX9* gene, which is not mutated in *ga2ox heptuple* mutant, and shows high expression levels in the root^24^, suggesting it may fulfil the role of the other *GA2OX* genes in *ga2ox heptuple* mutant. However, in our dataset, *GA2OX9* was not auxin responsive. Another explanation could be that the *ga2ox heptuple* mutant compensates for the absence of *GA2OX* genes by altering the expression of other genes involved in GA metabolism, as already shown in the *ga2ox quintuple* mutant^25^. Since all pathways responsible for GA homeostasis can be targeted to maintain optimal GA levels or response, to test this it would be necessary to analyse the expression of GA metabolic genes in the *ga2ox heptuple* mutant treated with auxin. The observed auxin-GA crosstalk via *GA2OX6* and *GA2OX8* might be significant only under specific environmental or developmental conditions, which we did not manage to test in our experiments. Auxin transport from the aboveground parts, regulated by environmental changes, could influence root elongation by modulating GA levels. Our findings align with this, showing that auxin transported from the shoots controls the expression of the GA2OX enzymes in the roots.

Additionally, several studies pointed to time-dependent GA response in roots. GA are important in long-term formation of lateral roots^44^. In contrast, previously we showed increased sensitivity of roots overexpressing dominant negative *AXR3-1* shortly after altering auxin signalling. Interestingly, these changes are not accompanied by measurable modulations in GA levels^3^. This could be explained by increased sensitivity of roots to GA. The positive effect of elevated GA sensitivity on root growth is also demonstrated by the increased root elongation of mutants overexpressing the GA receptor^45^.

Remarkably, although the GFP-RGA marker line^12^ indicated GA signalling decrease in response to auxin both in the meristematic and elongation zones, the GA level as determined by the direct nlsGPS2 sensor^36^ decreased in the elongation zone only. This can be explained by higher sensitivity of the GFP-RGA signalling reporter, perhaps via the high affinity and root expressed GID1B receptor, compared to the direct nlsGPS2 sensor that uses GID1C receptor action. Another explanation would be that auxin upregulates expression of RGA gene, however in our dataset, RGA was not auxin responsive.

High auxin levels in the stem cell niche (SCN)^39^ overlap with high *GA2OX8* expression, suggesting that auxin induces *GA2OX8* expression in these cells. Increasing GA level by pharmacological or genetical means positively correlates with the size of meristem and proliferation rate^46^. However, neither GA depletion or application affect the expression of genes important for SCN specification^46^, therefore more probable is that GA2OX8 is involved in controlling cell proliferation rate. Supporting this, the *ga2ox heptuple* mutant roots show increased productions of columella cell layers. GA biosynthesis genes are strongly expressed in QC^47^, *GA2OX8* might counteract their activity.

In our work, we have focused mostly on the root growth and root elongation zone. However, the observed strong *GA2OX8* expression in the root vascular system and its regulation by shoot-derived auxin suggests its role during vascular development. Mäkilä et al. showed^35^ that GA signalling regulates polar auxin transport to promote xylem production in root. GA2OX8 may catalyse the deactivation of GA12 – the most common precursor of bioactive GA present in vasculature^48^ as a part of feedback mechanism to balance vasculature formation.

Our results clearly demonstrate that auxin regulates GA level in the elongation zone through upregulation of *GA2OX6* and *GA2OX8* genes, and that GA level modulated by *GA2OX* genes is positively correlated with root cell elongation. However, numerous aspects of auxin-GA interactions in roots continue to be unresolved, with these processes affected by time, specific tissues, hormone concentration, environmental and developmental factors. Understanding the molecular mechanisms underlying auxin-GA crosstalk helps to understand the complexities of root cell elongation.

## Materials and Methods

### Plant material

Following lines were used in this article: pRGA::GFP-RGA^12^, nlsGPS2^36^, *ga2ox6* (SM_3_1859), *ga2ox8* (SALKseq_040686.2), pG1090::XVE>>*AXR3-1*^49^, pG1090::XVE>>*ΔMP*^50^, DR5::LUC^51^, *pGA2OX6::Venus-GA2OX6*^38^.

The following lines were prepared in this study. Above mentioned *ga2ox6* and *ga2ox8* were crossed to create *ga2ox6xga2ox8 double* mutant. To generate *ga2ox heptuple* homozygous mutant containing null alleles of seven *GA2OX*, *ga2ox1/2/3/4/6 quintuple* mutant^25^ was crossed with *ga2ox7-2xga2ox8 double* mutant (SALK_055721/SALKseq_040686). Subsequent generations were then backcrossed with *ga2ox1/2/3/4/6 quintuple* mutant until the homozygous *ga2ox heptuple* mutant was generated. *pGA2OX8::YFP-NLS* was crossed with pG1090::XVE>>*AXR3-1* and pG1090::XVE>>*ΔMP*. pRGA::GFP-RGA was crossed with pG1090::XVE>>*mCherry-GA2OX8*. *pGA2OX8::LUC*, *pGA2OX8::YFP-NLS*, pG1090::XVE>>*mCherry-GA2OX8* and *ga2ox heptuple* nlsGPS2 lines were prepared by molecular cloning as described below.

All lines used are in *Arabidopsis thaliana* ecotype Columbia-0 (Col-0) background. Primers used for genotyping are in Table S1.

### Growth conditions

Chlorine gas was used to surface-sterilize seeds^52^, followed by stratification for 2d at 4°C. Seedlings were grown vertically on plates containing 1% (w/v) agar (Duchefa) with ½ Murashige and Skoog (MS, Duchefa, 0,5g/l MES, 1% (w/v) sucrose, pH 5.8 adjusted with 1M KOH). Plants grown in growth chamber with 60% humidity, 22°C by day (16 h), 18°C by night (8 h), light intensity of 120μmol photons m^-2^s^-1^.

### Molecular cloning and plant transformation

We used GoldenBraid methodology^53^ as a cloning strategy. For stable transcriptional fusion of *GA2OX8* promoter, we cloned 1272bp upstream of *GA2OX8* (AT4G21200), including 12bp of its CDS and fused it with luciferase gene^54^ (*pGA2OX8::LUC* line) or YFP fused with NLS to C terminus^55,56^ (*pGA2OX8::YFP-NLS* line). Next, we cloned CDS of *AXR3-1* (AT1G04250) (88P→L substitution) or *ARF5/MP* (AT1G19850) driven by the 35S promoter and fused it either with mVenus^3^ (35S::*AXR3-1-mVenus*) or mScarlet-I^57^ (35S::*ARF5/MP-mScarlet*) to C terminus. All these constructs were terminated by 35S terminator and cloned into alpha1 vector.

For estradiol-inducible lines, XVE was cloned under the control of the G1090 promoter^58^ terminated by the RuBisCo terminator from *Pisum sativum* and cloned into alpha1-1 vector. We cloned CDS of *GA2OX8* downstream of the 4xLexA Operon driven by CaMV 35S minimal promoter^59^ and terminated by the 35S terminator, and we fused mCherry^60^ to the N-terminal part of *GA2OX8* CDS and terminated by 35S terminator. This construct was cloned into the alpha1-3 vectors^61^. The alpha transcriptional units were then interspaced with matrix attachment regions^61^. Alpha vectors were combined with a Basta resistance cassette in aplha2 plasmid and cloned into the pDGB3omega1 binary vector^58^. All constructs were transformed into the Col-0 ecotype using the floral dip method^62^. Primers used for cloning are in Table S2. *ga2ox heptuple* nlsGPS2 was generated though the Agrobacterium floral dip method and transformants were selected on kanamycin resistance.

### Treatments and phenotyping

To treat the plants, 5d old plants were transferred to treatment-containing medium. Estradiol (Sigma, 20mM stock in DMSO, 2,5μM working concentration), Indole Acetic Acid (IAA, Sigma,10mM stock in 96% ethanol) or propidium iodide (PI, Biotium, 1M stock in water) were used for treatments. Working concentrations and treatment time for specific experiments are given in the legend of each figure.

Root growth rate was measured as distance between root tip positions in consecutive time frames. Root length was measured as the distance between the root base and root tip. The size of the meristematic and elongation zone was measured on roots stained with 2,5μM PI for 15min. The end of the elongation zone was set as the first cell with root hair. The size of the meristematic zone was measured as a distance from the quiescent centre to the last isodiametric cell. In auxin treated roots, this rule was not feasible. Therefore, the boundary for the meristem was estimated as the end of the root cap. To test the role of shoot-derived auxin, shoots were decapitated with razor. To add exogenous auxin on shoot, plants were put on agar plates and 20μl of IAA treatment were added to cover shoot tissue. To avoid spreading the treatment to roots, 1-2mm gap in the agar was made between the shoot and root. Root width was measured in the region of 7 endodermal cell from QC, lateral root cap was not included in measurement.

### Microscopy, image analysis and low-resolution imaging

Vertical stage^63^ Zeiss Axio Observer 7 coupled to a Yokogawa CSU-W1-T2 spinning disk unit with 50μM pinholes and equipped with a VSHOM1000 excitation light homogenizer (Visitron Systems) was used for high resolution imaging. VisiView software (Visitron Systems, v4.4.0.14) was used to acquire images. Plants were imaged with a Plan-Apochromat 10x/0.45 M27, Plan-Apochromat 20x/0.8 M27 and LD LCI Plan-Apochromat 40x/1.2 Imm objectives. We used 561 nm laser for mCherry, mScarlet and PI-stained samples (excitation 561, emission 582-636 nm), 515 nm laser for mVenus, Venus and GFP (excitation 515, emission 520-570 nm) and 488 nm laser for YFP samples (excitation 488, emission 500-550 nm). Zeiss LSM 880 was used for PI stained roots, nlsGPS2 sensor and *pGA2OX8::YFP-NLS* crossed with *AXR3-1* or *ΔMP*. Roots were imaged by LD LCI Plan-Apochromat 25x/0.8 DIC Corr and C-Apochromat 40x/1.2Imm Corr DIC objectives. The GPS2 emission ratio of the nlsGPS2 sensor was calculated as the ratio of emission intensities of the Aphrodite-FRET acceptor channel (excitation 458 nm, emission 525-579 nm) and the Cerulean donor channel (excitation 458, emission 463-517 nm). In parallel, the Aphrodite channel was acquired (excitation 514, emission 525-579 nm). In the *pGA2OX8::YFP-NLS* crossed with pG1090::XVE>>*AXR3-1* or pG1090::XVE>>*ΔMP*, GFP and YFP signals were separated using lambda imaging and linear unmixing in the Zen software. Intensity of all fluorescent proteins were measured after removing the background intensity only from the nuclei.

Vertically placed flatbed scanner (Perfection V700, Epson) was used for low resolution imaging. Epson Scanner software v3.9.2.1US was used to acquire images. ImageJ Fiji software^64^ was used for all image analysis.

### Transcriptomic analysis and gene expression analysis

pG1090::XVE>>*AXR3-1*, pG1090::XVE>>*ΔMP* and Col-0 were grown on ½MS plates for 5d and then whole plants were poured with 5ml of 2,5μM estradiol-containing ½MS liquid medium, making sure that all plants are treated. Additionally, pG1090::XVE>>*AXR3-1*, pG1090::XVE>>*ΔMP* and Col-0 were poured with dmso-containing ½MS liquid medium. Leftover medium was removed, and plants grown vertically for 1.5h or 3h in cultivation room. After this, roots were harvested. To obtain transcriptomic data, RNA from 40-50 root tips (7-10mm) of pG1090::XVE>>*AXR3-1*, pG1090::XVE>>*ΔMP* and Col-0 was extracted following the protocol (Plant Total RNA Mini Kit, Favorgen). cDNA library was prepared from polyA-enriched total RNA and sequenced by Illumina. The sequencing resulted in at least 20 million 150 bps long read pairs. Rough reads were quality-filtered using Rcorrector and Trim Galore scripts ^65^. Levels of gene expression (aggregated transcript abundances quantified as transcripts per million – TPM) were determined using Salmon^66^ with parameters --posBias, --seqBias, --gcBias, --numBootstraps 30. Reference index was built from *Arabidopsis thaliana*, TAIR10 cds library, version 20101214. Visualization, quality control of data analysis and determination of differentially expressed genes were selected using sleuth (version 0.29.0) package in R^67^. Genes with q-value <= 0.05 and log2 fold change ≥ 1 (upregulated) or ≤ -1 (downregulated) were considered to be significantly differentially expressed.

### Luciferase reporter assay

We used *pGA2OX8::LUC*, p35S::*AXR3-1-mVenus* and p35S::*MP-mScarlet* in alpha1 vector prepared as described earlier. Constructs were transformed into *Agrobacterium tumefaciens* (GV3101), grown shaking overnight in LB medium at 28°C. Next day, cultured were centrifuged, and resuspended in infiltration buffer (10mM MgCl2 and freshly prepared 100µM acetosyringone) at OD600 = 0.3. An equal amount of cultures of different constructs were mixed for infiltration and incubated for 2h in dark at room temperature. 2ml of mixtures were injected into 4 weeks old *Nicatiana benthamiana* leaves. The activity of luciferase was measured 3d after infiltration using Azure 600 detector (Azure Biosystems). Chemiluminescence was measured on leaves or *pGA2OX8::LUC* in Arabidopsis transformants sprayed with 1mM luciferin, kept for 2min in dark, followed by 6min exposure.

### Statistical and graphical analysis

To compare multiple samples with normal distributed data, one-way ANOVA followed by Tukey HSD was used. Kruskal–Wallis test followed by post hoc Dunn’s test was used for data which do not follow normal distribution. Statistical analysis between two normally distributed samples was performed using the unpaired Student’s t-test (P-value 0.05 > ns; *P ≤ 0.05; **P ≤ 0.01; ***P ≤ 0.001).

Figures were generated using Inscape. Boxplots were made by SuperPlotsOfData^68^. Dots represent individual data points; the largest dot represents mean. Data analysis and line graphs were made by Microsoft Excel.

## Data availability

Rough sequencing data and quantification tables are available through GEO Series accession number GSE282145 at the Gene Expression Omnibus (https://www.ncbi.nlm.nih.gov/geo/).

## Acknowledgments

MF and MK received support from the European Research Council (Grant No. 101125499). MK was supported by Charles University Grant Agency (Grant No. 337021). MF and KM were supported from the project TowArds Next GENeration Crops, reg. no. CZ.02.01.01/00/22_008/0004581 of the ERDF Programme Johannes Amos Comenius. JG and AMJ were supported by the Gatsby Charitable trust (GAT3395) and European Research Council under the European Union’s Horizon 2020 research and innovation program (759282). The authors are grateful to Matouš Glanc for critical reading of the manuscript, Eva Medvecká for lab support and Professor George Coupland and Achard Patrick for kindly providing us with *pGA2OX6::mVenus-GA2OX6*, *ga2ox6* and pRGA::RGA-GFP seeds. Confocal laser scanning microscopy performed in the Microscopy Core Facility of Faculty of Science was co-financed by the Czech-BioImaging large RI project LM2023050.

## Supplemental data

Supplemental figure S1.

Supplemental Table 1. List of primers used for genotyping

Supplemental Table 2. List of primers used for cloning.

Dataset1 Excel file - Transcriptomic data

