## supplemental information for "Gibberellin-deactivating GA2OX enzymes act as a hub for auxin-gibberellin crosstalk in *Arabidopsis thaliana* root growth regulation"

A

| Promoter of gene | Position | Strand | Hit Sequence |
|---|---|---|---|
| **GA2OX8** (1272bp upstream) | 1238 | + | TGTCTc |
| **GA2OX6** (1750 bp upstream) | 164 | + | atGTCGGcta |
| | 163 | - | aaTGTCGgc |
| | 206 | + | TGTCTc |
| | 162 | - | taaTGTCGgc |
| | 163 | - | aaTGTCGgct |

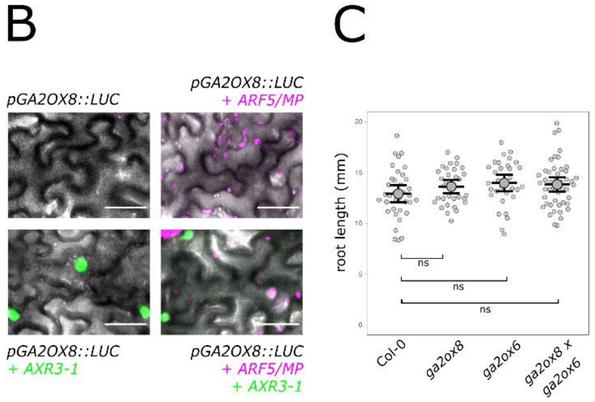


**Figure S1**

A. AuxRE binding sites in *GA2OX8* and *GA2OX6* promoter (obtained by PlantPAN 4.0)

B. Tabacco leaf cells co-infiltrated *pGA2OX8::LUC* with/without p35S::*ARF5-mScarlet* and p35S::*AXR3-1-mVenus*. Scale bar = 50μm.

C. Root length of 5d old Col-0, *ga2ox8*, *ga2ox6* and *ga2ox8xga2ox6 double* mutant.

The asterisks indicate statistically significant differences (ns – not significant, *P < 0.05, **P < 0.01, ***P < 0.001). Error bars in boxplots are CI.

### Table S1: List of primers used for genotyping

| Primer name | 5´-3´ primers | Lines |
|---|---|---|
| ga2ox1-1_RP | TCTTCCGGTTCGATATCTCCCA | *ga2ox septuple* mutant |
| ga2ox1-1_LP | TGACCAAAACACGGACTCGAT | |
| ga2ox2-1_RP | CTGCGAGGAGTTCGGGTTCTT | |
| ga2ox2-1_LP | TTTTTGTCGACCCTCCACACC | |
| ga2ox3-1_RP | TTGAAAATTTGTCCTTTAACCCCA | |
| ga2ox3-1_LP | TCCGGATGTGAAAACTGAAATCAA | |
| ga2ox4-1_RP | TGACAGCTCGGCAGTGAATTG | |
| ga2ox4-1_LP | TGGGGTATCACATTTCACCTCAA | |
| ga2ox6_2_RP | TTGTCAACCGTATGGAAACCG | |
| ga2ox6_2_LP | CAACCAAGAACCAACGATTGC | |
| ga2ox7-2_RP | AGTCACCATGGACTTTCGC | |
| ga2ox7-2_LP | GAGAAGTGGCGCTAGGGTTT | |
| ga2ox8_RP | AACGTTCGCAGACTCGTAGT | |
| ga2ox8_LP | ACCTTTCTTTTGGTTAAGTTACCTT | |
| ga2ox8_LP | TCGTACCTAGCTTGTTTTTAGAGG | *ga2ox6 x ga2ox8* double mutant |
| ga2ox8_RP | TGTGGAGAATTATCCCAATAACAAG | |
| ga2ox6_LP | TTGTCAACCGTATGGAAACCG | |
| ga2ox6_RP | CAACCAAGAACCAACGATTGC | |
| SLAT 3' dspm1_LP | CTTATTTCAGTAAGAGTGTGGGGTTTTGG | For ga2ox6 |
| WiscDs-Lox p745_LP | AACGTCCGCAATGTGTTATTAAGTTGTC | For ga2ox1 |
| SALK LB1.3_LP | ATTTTGCCGATTTCGGAAC | For ga2ox2/3/4/7 |

### Table S2: List of primers used for cloning.

| part | 5´-3´ primers used for domestication |
|---|---|
| pGA2OX8 | F: GCGCCGTCTCGCTCGGGAGAAGGAACAAAGCAACATTTGGATA<br>R: GCGCCGTCTCGCTCACATTAAAATACGTGTTGTGAGGAGAG |
| CDS GA2OX8 | F: GCGCCGTCTCGCTCGTTCGATGGATCCACCATTCAACGAA<br>R: GCGCCGTCTCGCTCAAAGCTTAGTAGACGTGATTAAGGAACC |
| CDS AXR3-1 | Part 1<br>F: GCGCCGTCTCGCTCGAATGATGGGCAGTGTCGAGCT<br>R: GCGCCGTCTCGCTGTCTCTGAGAACCCTCTC<br>part 2<br>F: GCGCCGTCTCGACAGTTGATCTGAAGCTAAATCTG<br>R: GCGCCGTCTCGTAAGACGTAAACGCTTGCATG<br>part 3<br>F: GCGCCGTCTCGCTTATGAAAGGATCGGATGCCATTGGTCTTGCTCCGAGG<br>R: GCGCCGTCTCGCTGTCTCTGAGAACCCTCTC |
| CDS MP | Part 1<br>F: GCGCCGTCTCGCTCGAATGATGGCTTCATTGTCTTGTG<br>R: GCGCCGTCTCGGGGACCCGCATATCGCCTTA<br>part 2<br>F: GCGCCGTCTCGTCCCAGCTCTCAGTTGGTAT<br>R: GCGCCGTCTCGTAAGACCGTTCAACTGAGTGT<br>part 3<br>F: GCGCCGTCTCGCTTAAGTTTGACCAGTTCAGTC<br>R: GCGCCGTCTCGCTCACGAACCTGAAACAGAAGTCTTAAGATCGTT |
